# Efficacy of the MEK1/2 inhibitor trametinib in combination with clinically-investigated γ-secretase inhibitors in rhabdomyosarcoma

**DOI:** 10.1101/622522

**Authors:** Megan M. Cleary, Douglas S. Hawkins, Charles Keller

## Abstract

The childhood muscle cancer rhabdomyosarcoma (RMS) is the most common pediatric soft tissue sarcoma. In the last 40 years, outcomes for low and intermediate risk patients have improved; however, high risk patients with metastatic disease still have poor overall survival. Differentiation therapy for RMS has been considered a potential clinical approach to halting tumor progression by inducing the terminal myogenic differentiation program, and thus reducing the need for cytotoxic chemotherapy. Both the NOTCH and MEK pathway have been shown to play varying roles in inducing differentiation in RMS cells. Here, we tested several different RMS cell lines harboring varying genetic abnormalities with the MEK inhibitor trametinib alone, and in combination with γ-secretase inhibitors and found no significant effect on cell viability when used together.

## Introduction

Rhabdomyosarcoma (RMS) is a rare pediatric cancer thought to phenocopy a skeletal muscle lineage. RMS presents as one of two main subtypes, each exhibiting a unique histological and molecular profile. Embryonal rhabdomyosarcoma (eRMS) is characterized by widespread genetic instability with *P53, RAS, PIK3CA, RB1* and FGFR4 disruptions often observed (1–6). Alveolar rhabdomyosarcoma (aRMS) is defined by a t(2;13) or t(1;13) chromosomal translocation that results in the DNA binding domain of either PAX7 or PAX3 fusing with the transactivation domain of FOXO1, creating an oncogenic transcription factor (7,8). Even though RMS is the most common soft tissue sarcoma in children, survival for patients with metastasis has remained unchanged over the past 47 years despite intensive multimodal therapy.

Histologically, rhabdomyosarcoma expresses markers of myogenic differentiation such as myogenin and MyoD1 (9,10); however, the function of these myogenic proteins is often impaired and RMS cells fail to fully differentiate (11). RMS is believed to circumvent terminal differentiation, allowing RMS tumor cells to divide uncontrollably. Restoring the terminal differentiation program is posited to slow or halt tumor growth by transforming malignant, proliferating cells into non-dividing cells. Differentiation therapy appears to have clinical potential, as eRMS cells have been observed to differentiate following chemotherapy and radiation (12–14), although this response is not often found in aRMS.

Different groups have uncovered strategies for inducing differentiation in RMS and thus slowing growth, but with limited success. Genetic suppression of the Notch1-Hey1 pathway by shRNA in eRMS RD cells, or suppression of Notch-3 by siRNA in RD and aRMS Rh30 cells results in an increase of myogenic differentiation markers *in vitro*, and pharmacological inhibition of Notch signaling using a γ-secretase inhibitor reduces cell proliferation (15,16). In RD cells, modulating *miR-206* increases differentiation by 30% *in vitro* (17), while GSK3 inhibitors significantly increase the number of myosin heavy chain positive cells after treatment for 72 hours (18). The inhibition of RAF/MEK protein kinases induce terminal differentiation in RD cells (18), a result not surprising given that RAS pathway activation is common in eRMS, and that this disease demonstrates a “Ras on” gene signature (5,19). *In vivo* trametinib slows but does not halt tumor growth in eRMS cell lines SMS-CTR and BIRCH, but not RD; these tumors exhibit an increase in nuclear MYOG expression, but not across all cells of a tumor, and complete terminal differentiation is not observed (20). Taken together, these results demonstrate the difficulty in achieving complete terminal differentiation and suggest the therapeutic benefit of a single agent differentiation therapy will have limited clinical success, and thus combining pathway inhibitors will likely be necessary to achieve tumor remission.

Unpublished data from our lab suggests *in vitro* synergy when simultaneously targeting the MAPK and Notch pathways in RMS cells. Presented here are the *in vitro* drug screening assays performed on a range of aRMS and eRMS cells lines and primary patient cells harboring different genetic hallmarks, to examine the efficacy of the MEK inhibitor trametinib alone or in combination with Notch signaling inhibition using γ-secretase inhibitors.

## Results

### Trametinib decreases viability in a range of cell lines

Our experiments confirmed that treatment of KRAS-driven RD cells with sub-micromolar concentrations (684 nM) of trametinib for 72 hours resulted in cytotoxicity (Figure 1A). Unexpectedly, SCA1-01, a Kras activated primary mouse tumor cell line, was not sensitive to MEK inhibition (IC50= 38,000 nM), indicating that a pathway other than RAS-MEK is necessary for tumor cell maintenance (Figure 3). An aRMS patient-derived primary tumor cell culture (CF-1, Figure 2A) and an aRMS cell line (Rh30, Figure 4A) that both harbored the PAX3:FOXO1 chromosomal translocation were also treated with trametinib and responded with low micromolar concentrations (IC50= 1,557 nM and 803 nM, respectively). Finally, we tested CW9019, a cell line that has an alveolar histology and harbored a t(1;13) reciprocal translocation resulting in a PAX7:FOXO1 fusion protein. This cell line was the most sensitive to trametinib treatment, with a half maximal inhibitory concentration of 113 nM (Figure 5A). However, the *in vitro* IC50 values for these cell lines are still well above the clinically achievable dose of trametinib (36 nM) (21).

**Fig 1.**
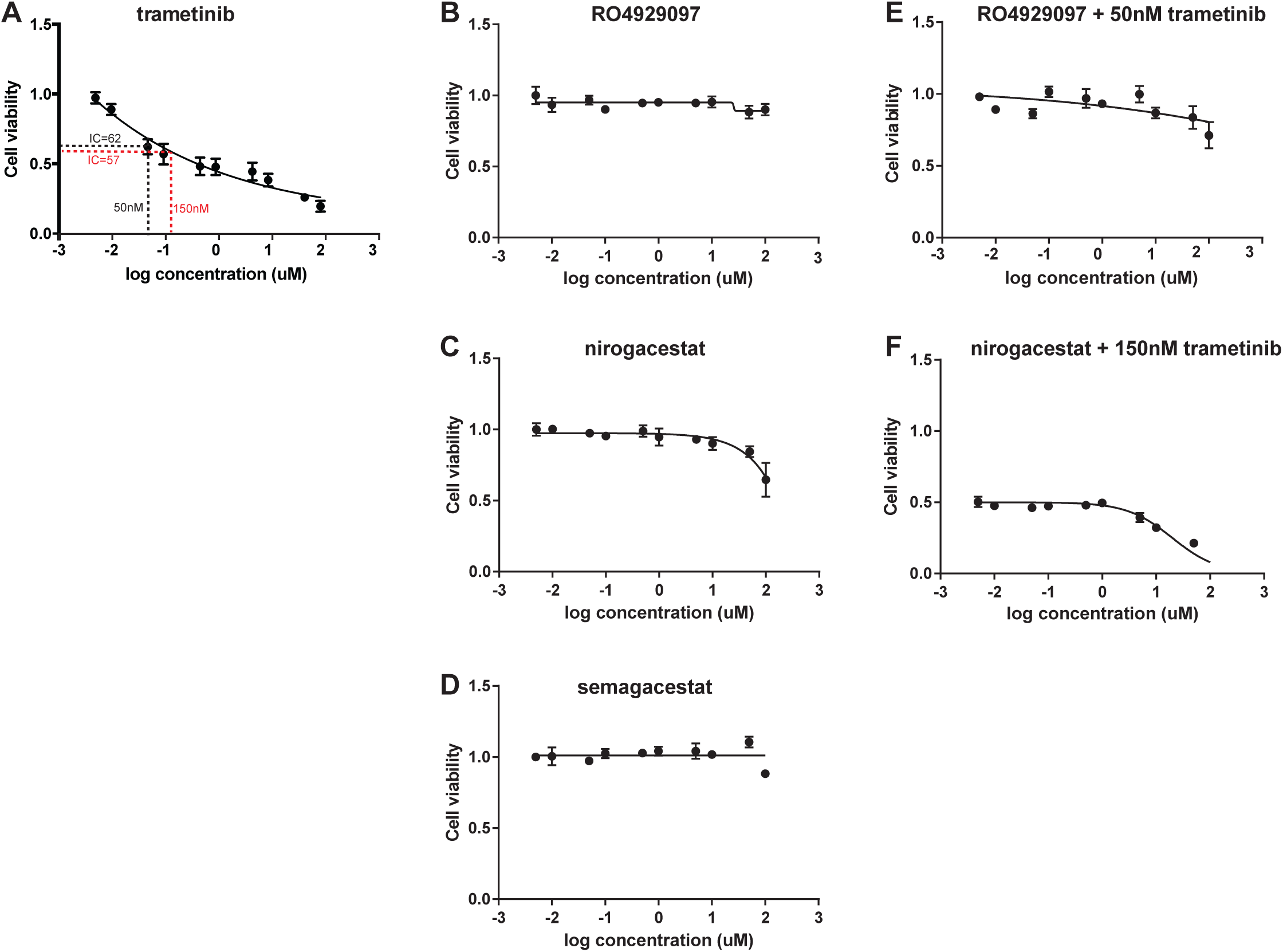
Cell viability assays using the KRAS-driven human eRMS cell line RD. Cells were treated with trametinib (A) or γ-secretase inhibitors (B-D) alone, or 50 nM (IC62) trametinib (E) or 150 nM (IC57) trametinib (F) in combination with varying concentration of γ-secretase inhibitors.

**Fig 2.**
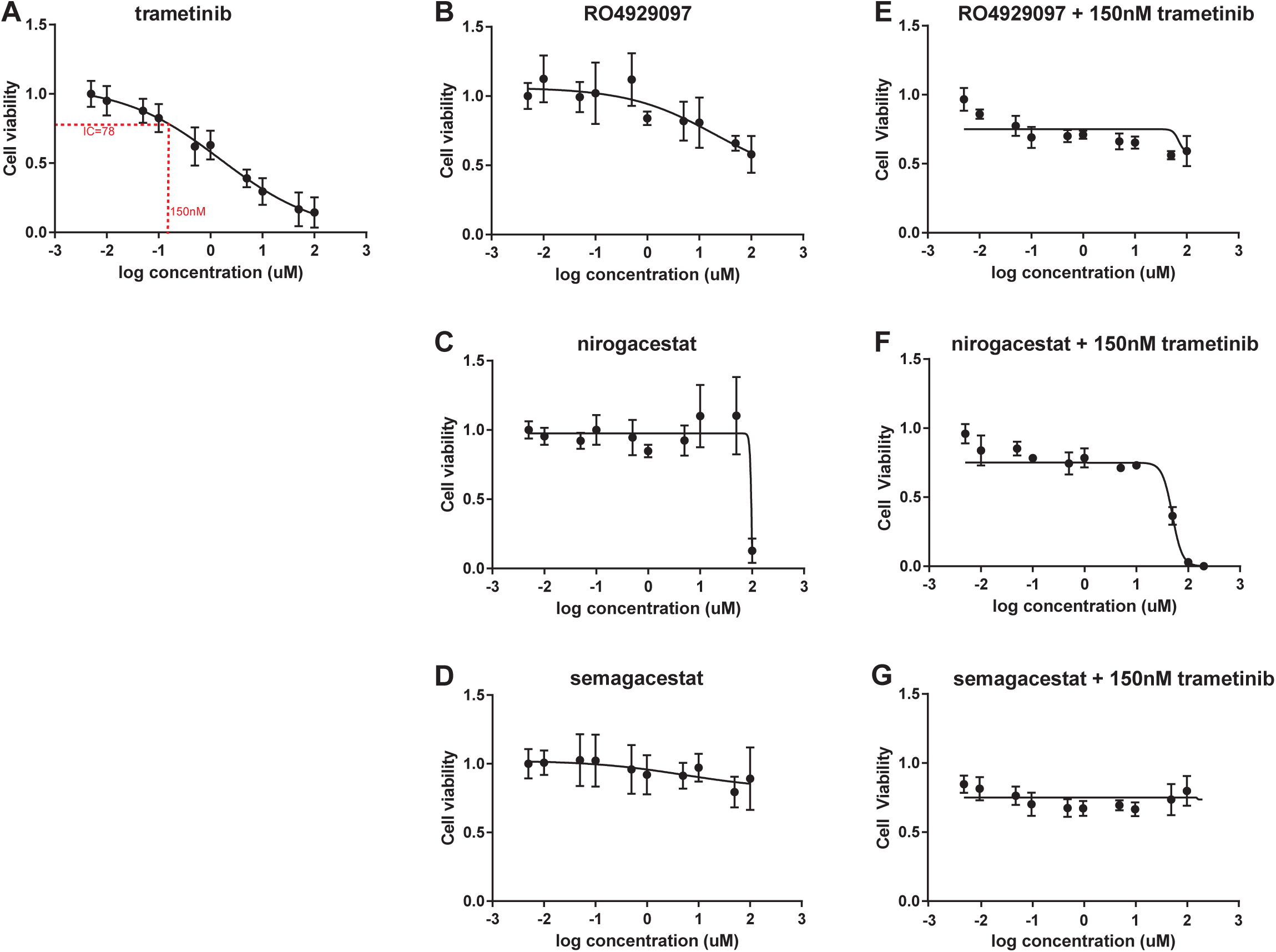
Cell viability assays using the primary patient derived aRMS cell culture CF-1. Cells were treated with trametinib (A) or γ-secretase inhibitors alone (B-D), or 150 nM (IC78) trametinib in combination with varying concentration of γ-secretase inhibitors (E-G).

**Fig 3.**
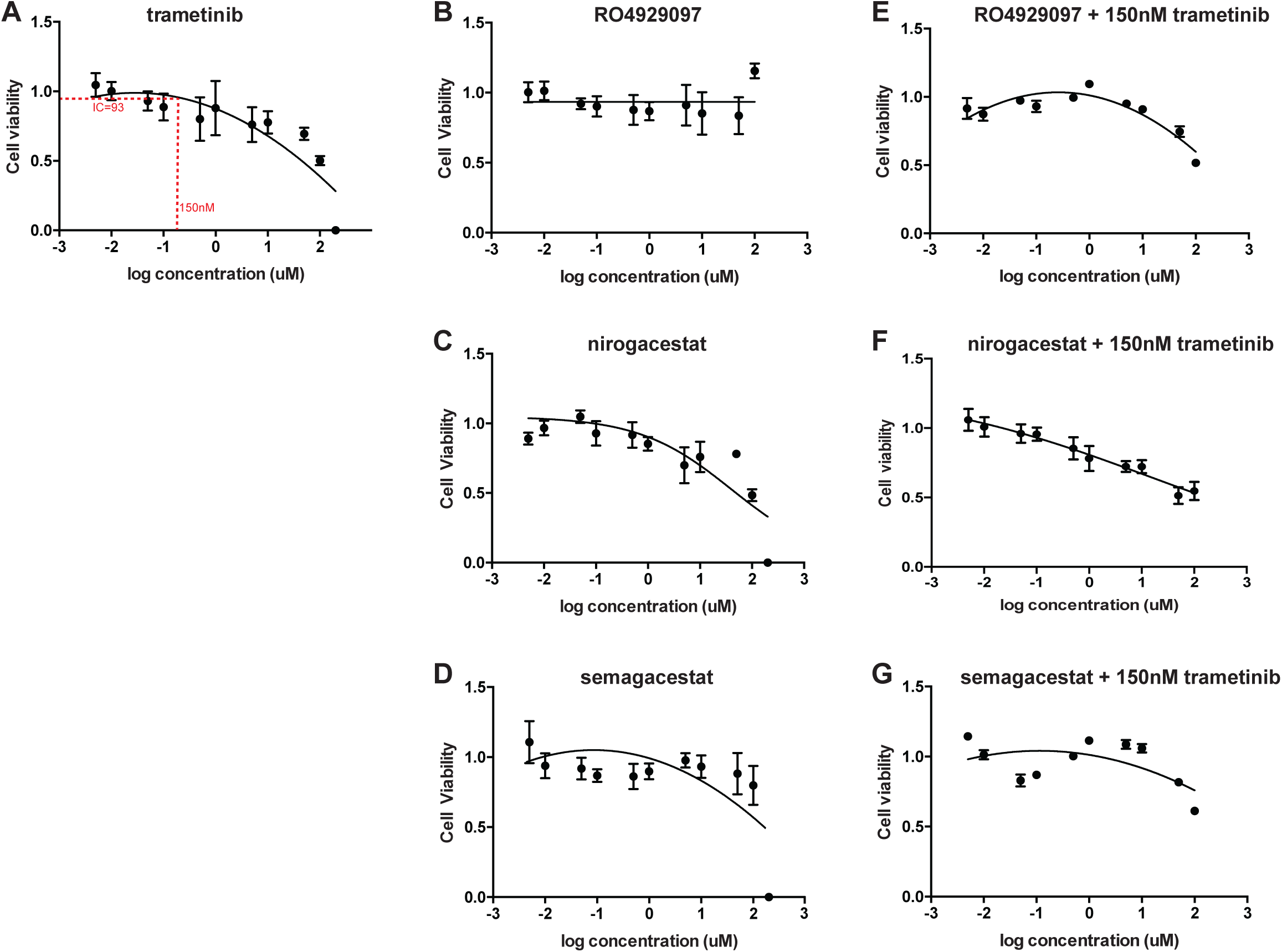
Cell viability assays using the *KRAS;p16p19^null^ (Cdkn2a)* mouse non-myogenic soft tissue sarcoma tumor SCA-1. Cells were treated with trametinib (A) or γ-secretase inhibitors alone (B-D), or 150 nM (IC93) trametinib in combination with varying concentration of γ-secretase inhibitors (E-G).

**Fig 4.**
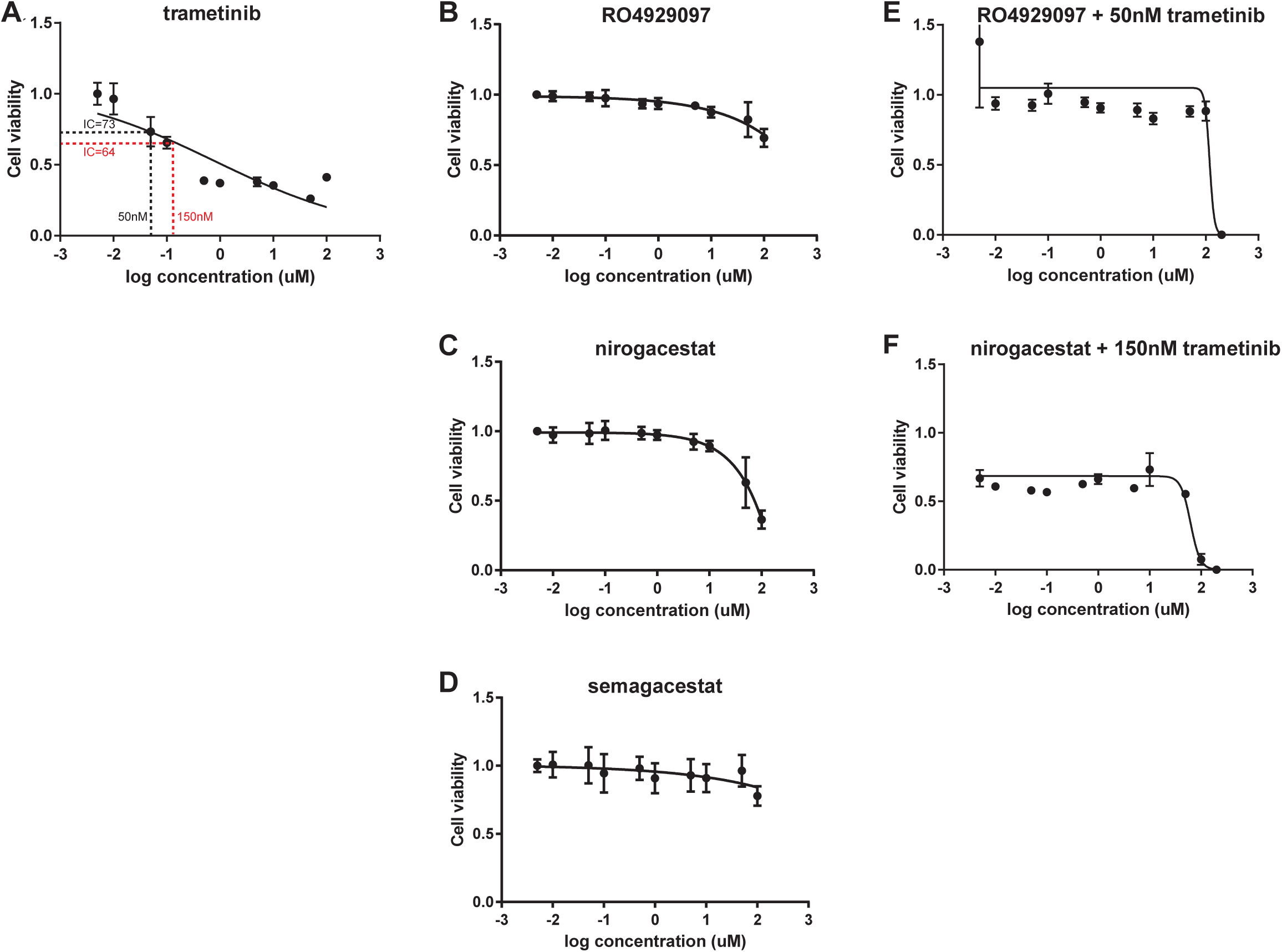
Cell viability assays using the human aRMS cell line Rh30. Cells were treated with trametinib (A) or γ-secretase inhibitors alone (B-D), or 150nM (IC64) trametinib (F) or 50 nM (IC73) trametinib (E) in combination with varying concentration of γ-secretase inhibitors.

**Fig 5.**
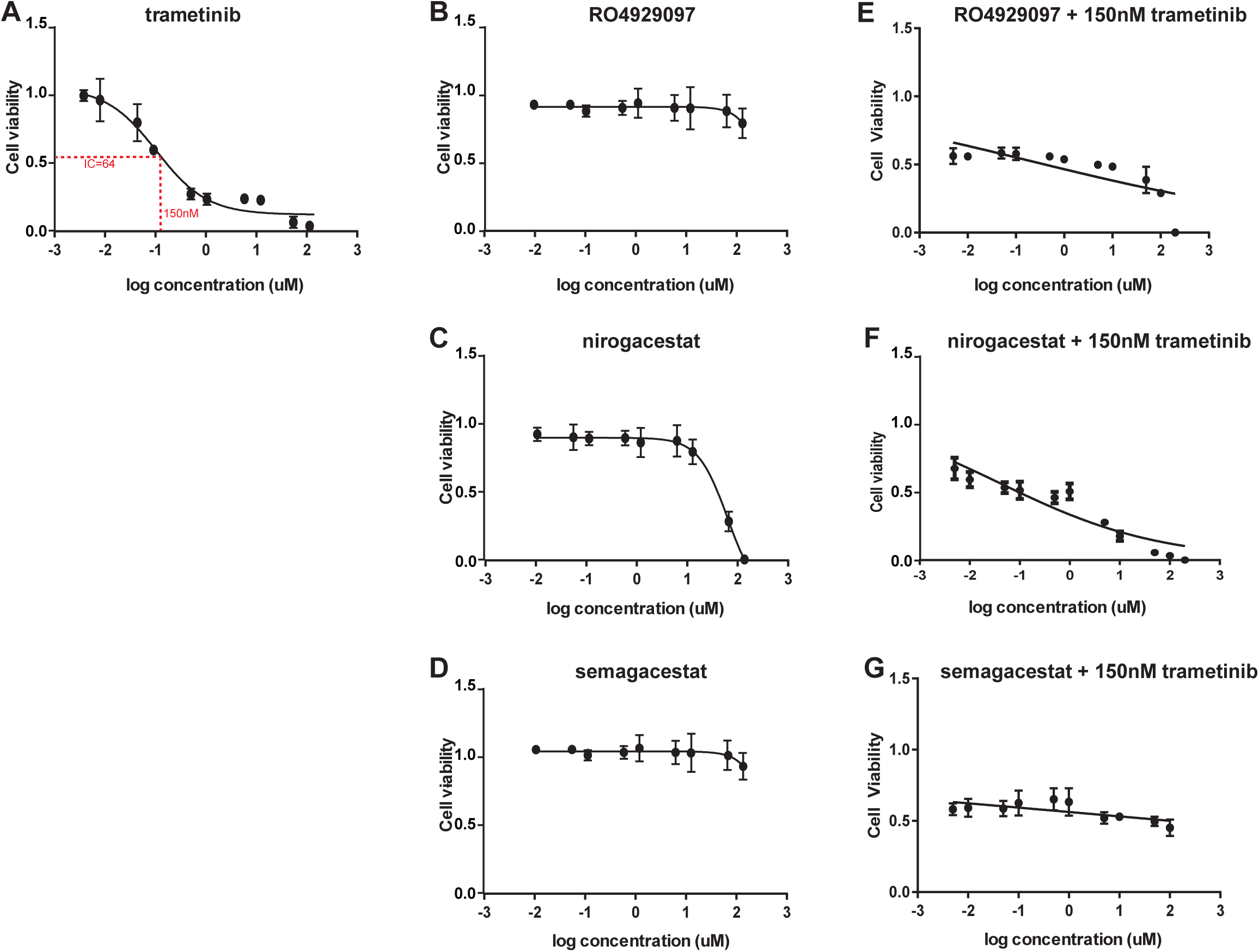
Cell viability assays using the human aRMS cell line CW9019. Cells were treated with trametinib (A) or γ-secretase inhibitors alone (B-D), or 150nM (IC64) trametinib in combination with varying concentration of γ-secretase inhibitors (E-G).

### Trametinib in combination with γ-secretase inhibitors does not exhibit synergistic cytotoxicity

To query whether trametinib augmented with inhibition of the Notch pathway would cause additive or synergistic cytotoxicity, we additionally tested the above cell lines in combination with one of the clinical γ-secretase inhibitors, nirogacestat, semagacestat or RO4929097. γ-secretase inhibitors were combined with either a low, clinically achievable dose (50 nM) or a high dose (150 nM) of trametinib. Little or no synergy was observed at clinically achievable concentrations of γ-secretase inhibitors in all cell lines tested. In RD cells γ-secretase treatment alone did not cause cytotoxicity (Figure 1B-D); synergy was only observed at the highest concentrations of nirogacestat and the highest tested concentration of trametinib (150nM, Figure 1F), but not with RO4929097 and a low concentration of trametinib (50nM, Figure 1E). γ-secretase inhibitors exhibited no synergistic effect on the aRMS primary patient sample cell culture CF-1 (22), the aRMS cell line Rh30 or the mouse non-myogenic sarcoma tumor SCA-1 when combined with trametinib (Figure 2, 3, 4 E-F). The aRMS cell line CW9019 which was most sensitive to single agent trametinib treatment exhibited some single agent sensitivity to the highest concentrations of nirogacestat (Figure 5C) but not semagacestat or RO4929097 (Figure 5 B,D). Combination treatment with 150 nM trametinib and γ-secretase inhibitors did not demonstrate any synergistic decrease in cell viability (Figure 5 E-F).

## Discussion

A common assumption in clinical trial design is that single agents should have strong activity in phase I/II studies to advance to phase III trials. However, some drugs may have substantial synergy and might improve survival by 10-20% in Phase III clinical trials, even if no activity could be seen in phase I/II trials. In this context, we sought to test the combination of two drugs with limited single agent potential, MEK inhibitors and γ-secretase inhibitors. Unfortunately, even in the earliest *in vitro* pre-clinical studies, no supporting evidence for this combination approach could be garnered.

## Materials and methods

### Cell lines

SCA1-01 cells were donated by Dr. Amy Wagers and Dr. Simone Hettmer (Harvard University) and established from a Kras;p16p19^null^ mouse non-myogenic sarcoma tumor. CW9019 cells were a generous gift from Dr. Fred Barr (National Cancer Institute). SCA-1 and CW9019 were maintained in DMEM supplemented with 10% FBS and 1% penicillin-streptomycin and maintained at 37°. RD (CCL-136) eRMS and Rh30 (CRL-2061) aRMS cells were purchased from ATCC (Manassas, VA) and cultured per manufacturer’s instructions. CF-1 is a primary patient sample isolated from a 18 month old male presenting with alveolar rhabdomyosarcoma (22). These cells were cultured in RPMI supplemented with 10% FBS and 1% penicillin-streptomycin and maintained at 37°. CF-1 cells are only tested at passage earlier than 8. All human cell lines were authenticated using short tandem repeat analysis performed by the University of Arizona Genetics Core (Tuscon, AZ).

### *In Vitro* inhibitor testing

Small molecule inhibitors semagacestat (LY-450139, S1594), nirogacestat (PF03084014, S1575) and RO4929097 (S8018) were purchased from Selleckchem (Houston, TX), reconstituted to manufacturers specifications and stored in −80°C. To generate a standard 10-point dose response curve, inhibitors were distributed as single agents into 96 well plates to produce final concentrations ranging from 100 μM to 0.005 μM. Combination plates were likewise prepared in 96 well plates with trametinib (GSK1120212, S2673) at fixed final concentrations of 50nM or 150nM, paired with γ-secretase inhibitors at varying concentration identical to those used for single agents. All drug plates were stored at −20°C and thawed a maximum of 3 times. Cells were seeded at a density of 3×10^4^ cells per well in a 96 well plate on day 0 (0h). On day 1 (24h) each drug or combination was added to the cells for a final volume of 100μL per well. Cells were incubated at 37° with 5% CO_2_ for 72 hours. On day 4 (96h) compounds were screened for their effect on proliferation by adding 100μL room temperature cell titer-glo (CTG, Promega, Madison, WI) to each well and rocking in the dark at room temperature for 10 minutes. Luminescence was measured with a Biotek Synergy plate reader. Standard dose response curves and IC50 values were calculated with GraphPad Prism software using log-10 transformed concentrations and normalized values. Each dose was performed in triplicate and all experiments were repeated three times.

**Table 1.** Cell lines used for this study.

| Name | Origin | Genetic features | Histology |
| --- | --- | --- | --- |
| RD | 7 year old female | MYC amplification, NRAS Q61H mutation, TP53 mutation | eRMS |
| Rh30 | 16 year old male | t(2;13), PAX3:FOXO1, TP53 mutation, amplification of 12q13-15 region including *CDK4* | aRMS |
| CF-1 | 5 year old male | t(2;13), PAX3:FOXO1 | aRMS |
| CW9019 | human | t(2;13), PAX7:FOXO1, TP53 mutation | aRMS |
| SCA-1 | mouse | KRAS, p16p19 null | NMS (non-myogenic sarcoma) |

**Table 2.**
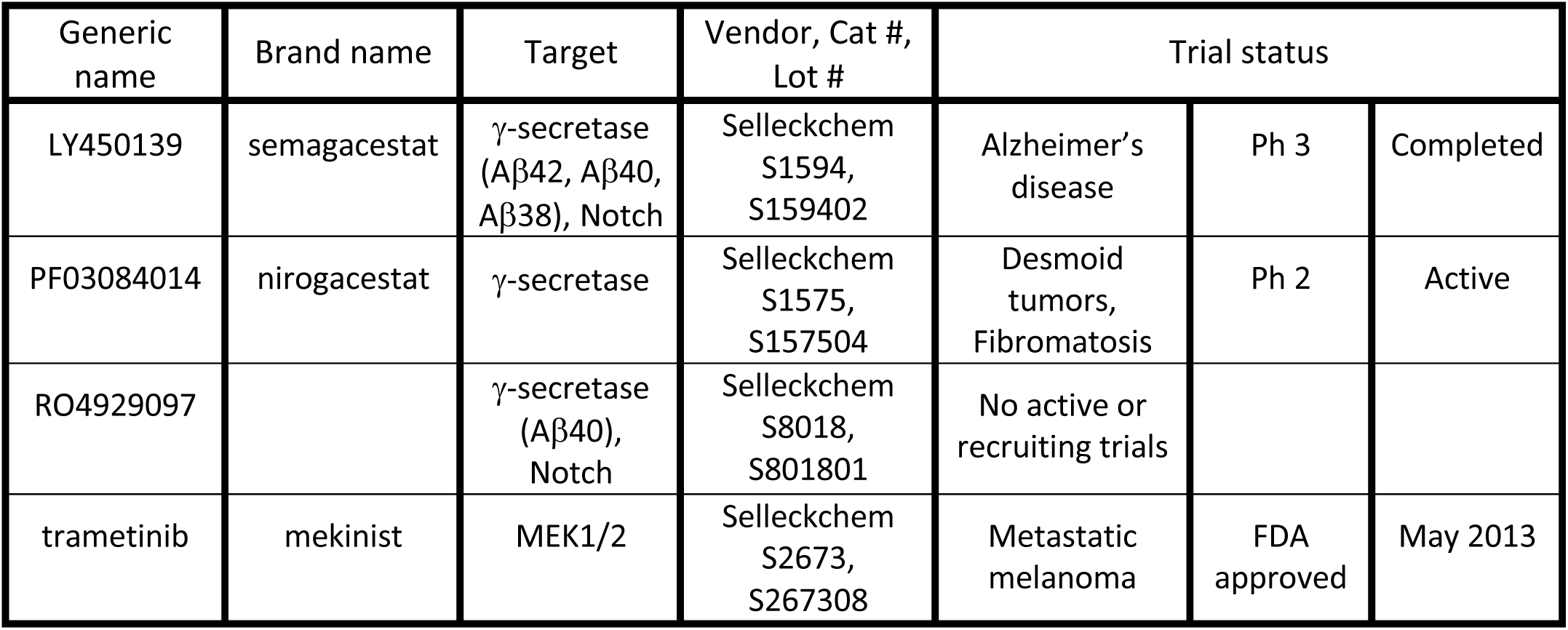
Drugs used for this study.

| Generic name | Brand name | Target | Vendor, Cat #, Lot # | Trial status |  |  |
| --- | --- | --- | --- | --- | --- | --- |
| LY450139 | semagacestat | γ-secretase (Aβ42, Aβ40, Aβ38), Notch | Selleckchem S1594, S159402 | Alzheimer's disease | Ph 3 | Completed |
| PF03084014 | nirogacestat | γ-secretase | Selleckchem S1575, S157504 | Desmoid tumors, Fibromatosis | Ph 2 | Active |
| RO4929097 | | γ-secretase (Aβ40), Notch | Selleckchem S8018, S801801 | No active or recruiting trials | | |
| trametinib | mekinist | MEK1/2 | Selleckchem S2673, S267308 | Metastatic melanoma | FDA approved | May 2013 |

## Acknowledgements

This work was supported in part by NCI R01 grant R01CA189299 as well as The Building Blocks Project.

We thank Dr. Amy Wagers and Dr. Simone Hettmer for donations of cell lines and Dr. Srinath Sampath and Dr. Sihari Sampath helpful discussions and advice.

## Competing Interests Statement

CK collaborates with Novartis in no-cost studies, and Novartis is the marketer of trametinib.

## Compliance with ethical standards

No human subjects or animals were used in these studies.

## Additional Information

### Data Deposition and Access

No genomic data sets apply to this study

### Author Contributions

MMC, CK, DSH participated in study design. MMC performed experiments. MMC, CK participated in data analysis. MMC, CK participated in writing manuscript.

